# Association Plots: Visualizing associations in high-dimensional correspondence analysis biplots

**DOI:** 10.1101/2020.10.23.352096

**Authors:** Elzbieta Gralinska, Martin Vingron

## Abstract

In molecular biology, just as in many other fields of science, data often come in the form of matrices or contingency tables with many measurements (rows) for a set of variables (columns). While projection methods like Principal Component Analysis or Correspondence Analysis can be applied for obtaining an overview of such data, in cases where the matrix is very large the associated loss of information upon projection into two or three dimensions may be dramatic. However, when the set of variables can be grouped into clusters, this opens up a new angle on the data. We focus on the question which measurements are associated to a cluster and distinguish it from other clusters. Correspondence Analysis employs a geometry geared towards answering this question. We exploit this feature in order to introduce *Association Plots* for visualizing cluster-specific measurements in complex data. Association Plots are two-dimensional, independent of the size of data matrix or cluster, and depict the measurements associated to a cluster of variables. We demonstrate our method first on a small data set and then on a genomic example comprising more than 10,000 conditions. We will show that Association Plots can clearly highlight those measurements which characterize a cluster of variables.

## 1. Introduction

A fundamental question in genomic data analysis is the following: Given a cluster of conditions (variables), which genes (measurements) are characteristically highly-expressed in these conditions, i.e. are associated to these conditions? Approaches to this question occur in many forms, be it as biclustering (Tanay *and others*, 2002; Pontes *and others*, 2015) or in the search for marker genes. While for small data sets answering this question is fairly easy, for data sets with a higher number of conditions the situation is more complex and poses a significant challenge to analysis and visualization methods currently available.

Although methods such as principal component analysis (PCA) have been successfully employed for many years, they have serious limitations when applied to large and complex data sets. For such data the first two or three principal components may explain only a small fraction of the total variance, often well below 5% of the total variance. This renders the low-dimensional representation, obtained after projection, effectively pointless due to the massive loss of information. We are faced with the challenge of visualizing information from higher dimensions.

In this work we focus on the analysis and visualization of a large data-set in the presence of a known clustering of conditions. This is a realistic setting since clustering has become a routine step in data analysis, or, in many cases, a clustering is imposed naturally by the structure of the data. Given a cluster, we will define an Association Plot, which can visualize the cluster-specific gene sets in a manner independent of the size of the dataset. Association Plots are derived from correspondence analysis (CA) (Greenacre, 1984), itself a data projection technique closely related to PCA. In CA the joint embedding of measurements (rows) and variables (columns) from a data matrix in one space, conveys information as to the association between variables and measurements. We take advantage of this CA property for our definition of Association Plots. The Association Plot is planar, although it is not a projection, and it stays two-dimensional even when representing very high-dimensional data.

This paper is organized as follows. First, we will introduce the formalism behind CA and Association Plots. A data example will serve to illustrate the new visualization. Subsequently, we will introduce SVD-based denoising and address some statistical and scoring issues about the visual patterns one observes. Lastly, we will apply the new methods to a very large gene expression dataset.

## 2. Contingency tables and association

We assume that data are given in the form of a matrix with non-negative entries. The methods we are going to use have traditionally been applied to contingency tables, which contain count data and are therefore also integer valued. For our purposes it suffices that the entries of the matrix are non-negative but not necessarily integer. In terms of application, we think, e.g., of gene expression values determined using the RNA-seq technology (Wang *and others*, 2009). A row corresponds to transcript abundances, while a column corresponds to the biological condition under which the measurement was performed. Like with any contingency table, we will look at row-frequencies and column-frequencies, and compare observed frequencies in a matrix cell with the cell’s expected frequency.

Our particular question pertains to the associations between rows and columns, or between rows and a cluster of columns. For a mathematical definition of association we follow the logic of contingency tables, where the likelihood ratio for the entry in cell (*i, j*) to be a product of chance would be 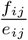, the observed frequency in the cell divided by the expected frequency. When this ratio is close to 1, we have little reason to believe that there is an association, whereas a large ratio hints at an association between the row and the column. Subtracting 1 from the ratio we can write this as 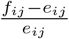, which will be near 0 in the absence of an association. We call this quantity the “association ratio” and abbreviate it as *a*(*i,j*).

Since we are also interested in the association between genes and a cluster of conditions, we proceed to add up the association ratio for the respective cells of the matrix. Let 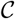, 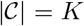 be a set of conditions *j*_1_,…, *j*_K_. We extend the notation to clusters by defining

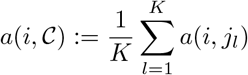

and call this the association ratio of the gene with the set of conditions. Of course, for the set of conditions we a have a cluster in mind, whose conditions behave similarly.

## 3. A rehash of correspondence analysis

Correspondence analysis (CA) (Greenacre, 1984, 2017) is a projection method for visually representing a data matrix in a high-dimensional space. Unlike PCA, CA does not submit the data matrix itself to a singular value decomposition, but the “object of interest” is the matrix of Pearson residuals derived from the data matrix. CA is computed in the following steps. Let *M* be a matrix with *G* rows and *C* columns. By *m_gc_* we denote a value in the *g^th^* row and *c^th^* column, and by *m*_++_ the grand total of *M*. One calculates an observed proportion matrix *P* with elements *p_gc_* = *m_gc_*/*m*_++_ and uses this for calculating row and column masses. The g^th^ row mass, *p*_*g*_+, is defined as the sum across row *g*, and the *c^th^* column mass, *p*_+*c*_, as the sum of all values in the *c^th^* column of *P*. With the expected proportions *e_gc_* = *p*_*g*+_ * *p*_+*c*_ Pearson residuals 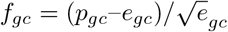. are computed. For comparison to contingency tables, note that the sum of their squares would form the χ^2^ statistic.

In analogy to PCA, it is now this matrix *F* = (*f_gc_*) that gets submitted to singular value decomposition, factoring it into the product of three matrices: *F* = *USV^T^*. *S* is a diagonal matrix and its diagonal elements *s_cc_* are known as singular values of *F*. Furthermore, the matrices *U* (with elements *u_gc_*) and *V* (with elements *v_gc_*) contain left- and right singular vectors, respectively, which are represented by the columns. From the left and right singular vectors coordinates of points for rows and columns in an high-dimensional space are computed. The coordinates *v* depicting row *g* are defined as 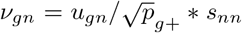, for *n* = 1,…, *c* (Greenacre, 2017, (A.8)). The coordinates *ω* representing column c are given by 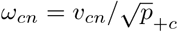 (Greenacre, 2017, (A.7)). In the literature (Greenacre, 2017) this choice of scaling is called ‘‘asymmetric”, with the rows represented in ‘principal coordinates” and the columns in ‘standard coordinates”.

It is a key feature of correspondence analysis that one can interpret the joint map of points for rows, the *v*’s, and columns, the *ω*’s, in the same space. The (full) dimension of this space will be *m*:= min(*C, G*)-1 (Greenacre, 2017, p. 203) and both sets of points can be thought of as elements of 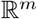. We will refer to this space as the CA-space. Traditionally, one uses only the first 2 or 3 dimensions and calls this a biplot. In the examples below we will, where meaningful, also depict the 2-dimensional biplot, although only for illustration purposes. The focus of our exposition, however, is on large data sets where too much information would be lost upon projection into 2 dimensions. Instead of ‘explained variance” that is used in PCA, CA speaks of “inertia” which gets approximated better with increasing number of dimensions used.

The geometry of the Association Plots to be defined below rests on two key features of CA (Greenacre, 2017):

1. A column-point can be expressed as a weighted sum of row-points, where the weights are related to the contribution of the rows to the particular column in *P*. This is Greenacre’s transition equation (Greenacre, 2017, (A.16)). It is the reason for the clustering of similar rows (genes) and similar conditions (columns), respectively, in CA space.
2. What is even more important for our application is that the inner product of row- and column-vector approximates the respective association ratio, i.e.

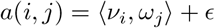

where in the m-dimensional CA-space *ϵ* = 0. This is Greenacre’s reconstitution formula (Greenacre, 2017, formula (13.5) or (A.14)) and it is a consequence of the SVD which was used to compute the coordinates. It pertains directly to our goal of describing association in geometrical terms: Due to the inner product, the row-vectors (genes) that are associated to a column-vector (condition) lie in the direction of that column; the more aligned the two are, and the longer the vectors, the higher the association. When a low-dimensional projection is permissible, this feature is usually clearly visible.

Note that the reconstitution formula also allows for generalization to clusters: row-points that are associated to a cluster of columns (conditions) lie in the direction of these (clustered) conditions. Since the inner product is bilinear, it also means that the association ratio sums nicely over groups of genes or clusters of conditions. This will be utilized below.

## 4. Association plots

A cluster of conditions 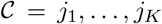 can be represented by the centroid of its condition vectors *ω_gi_*, *l* = 1… *K* in CA-space. We call this centroid 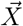:

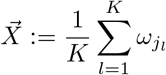

Due to linearity, we can see the average association ratio also in the inner product between a gene and the vector from the origin to this centroid. Let 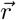 be a gene-vector in CA-space representing row *r* of the data. Then we can express the association between *r* and the cluster 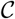 as:

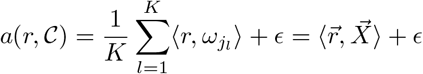

This is a trivial consequence of the reconstitution formula and the definitions from above.

This observation forms the basis for a simple 2-dimensional visualization of the high-dimensional information. The inner product is determined by the length of 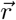, 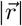 the length of 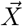, 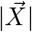, and the angle between the two vectors. We call the angle 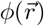, or just *ϕ* where the context is clear. In this notation, the inner product from above can be written as

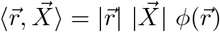

Therefore it makes sense to introduce a 2-dimensional representation where the x-axis corresponds to the direction of the centroid vector, and we represent 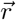 by the following x- and y-coordinates:

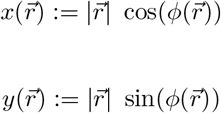

Clearly, 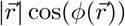 is the length of the projection of 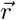 onto 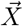, or 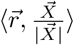, and 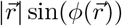 is the length of the orthogonal distance of 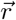 to 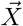, or 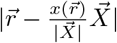. Also, 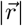 is equal to the length of the vector 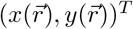. We define the Association Plot for cluster 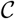 as the 2-dimensional plot where each gene-vector 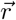 in CA-space is represented by these 2-dimensional points 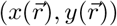.

Introducing 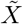 as the 2-dimensional vector

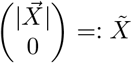

we can ascertain the conservation of the inner products, and with it the association ratio, between CA-space and Association Plot:

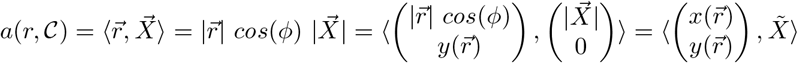

This demonstrates that the association ratio can be seen both as an inner product in the highdimensional CA-space as well as an inner product in the 2-dimensional Association Plot. For easier reading we have left out the error term *ϵ*.

From this simple line of algebra above one also notes that 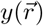 gets multiplied with 0. This means that the inner product, and with it the association ratio between gene and cluster, is constant along vertical lines in the Association Plot. Intuitively, this is due to the fact that by definition the association ratio 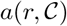 calculated for a given condition cluster is not influenced by other conditions or clusters in the data, which might also attract a gene. The angle *ϕ* contributes information because the less competition there is for a gene from other clusters, the smaller will be *ϕ*, whereas when a gene is shared also by other clusters, this will be reflected in a larger *ϕ*. This will be visible in the example below, and will be studied further in the section on significance of the visual patterns.

For easier interpretation one can also embed conditions into the Association Plot using the same coordinate system of projection length onto the centroid vector and orthogonal distance to it. As will be visible in the examples below, this conveys intuition about the coherence within a cluster.

## 5. A simple data example

Before continuing to develop Association Plots we present a simple, non-biological example in order to illustrate the concept of studying the row-column associations using Association Plots. The data set comes from the International Labour Organization and describes people’s sector of employment in 233 countries (International Labour Organization, 2020). Employment sectors are agricultural sector, industry sector, services sector, or unemployed. Data are further divided according to gender (m/f) and year (2000, 2005, 2010, 2015). This results in a matrix with 233 rows for the countries and 32 columns for employment category, gender, and year. For example, the column “industry, female, 2005” would represent the percentage of female population of a given country that was employed in industry in 2005.

The categories agriculture, industry, services, unemployed - possibly refined by gender - form natural clusters in the data. This is easily confirmed by visual inspection of, e.g., the 2-dimensional CA projection (Fig. 1). The year from which the data comes seems to have a lesser influence - at least in the 2D projection.

**Fig. 1:**
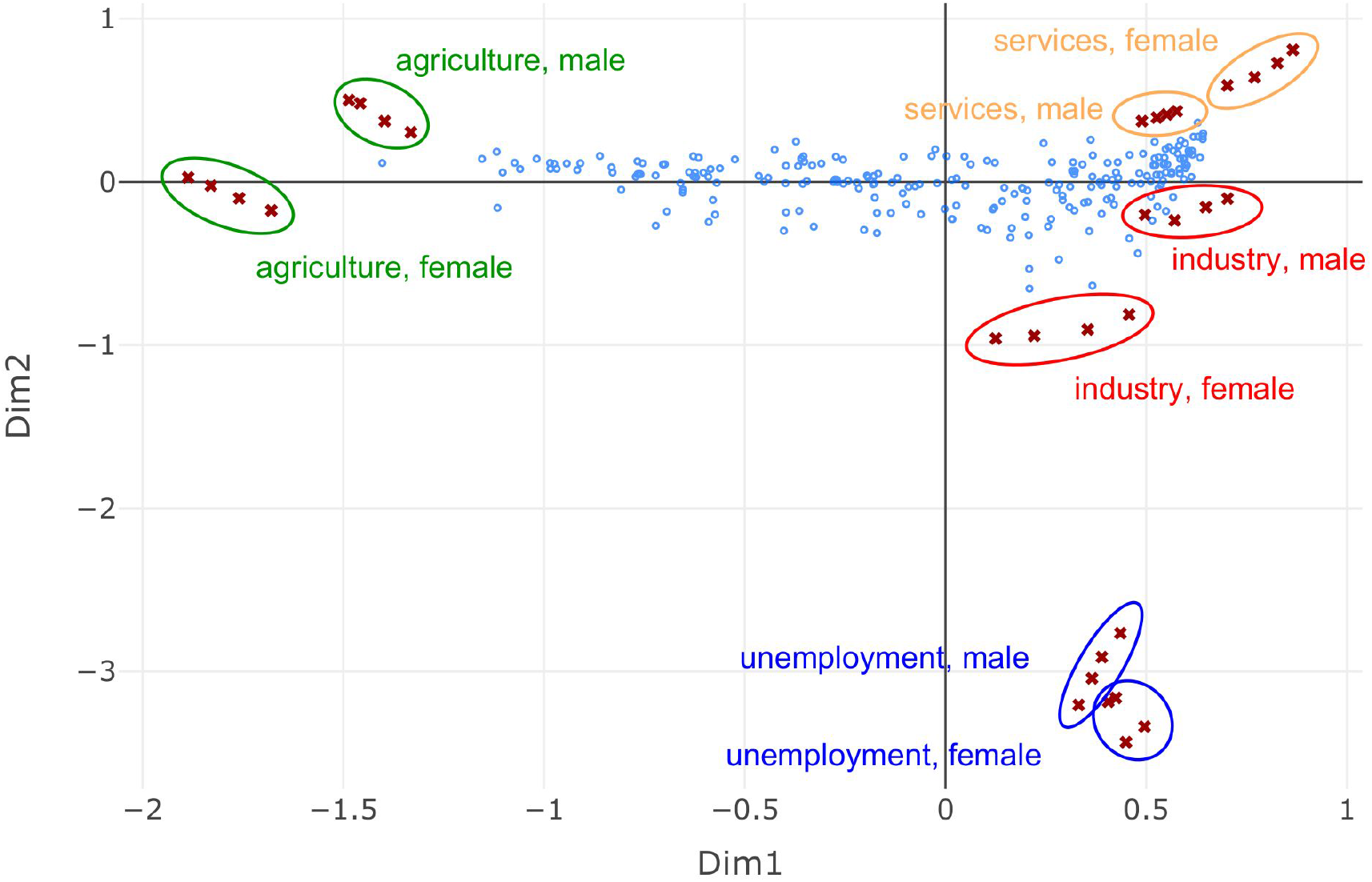
Two-dimensional CA projection of the employment data. The projection allows discerning four main clusters: agriculture sector, services sector, industry sector, and unemployed. Each of these clusters contains data points from four years (2000, 2005, 2010, and 2015). The location of the countries, represented in the biplot by blue dots, implies their employment profiles in the years 2000-2015.

The blue dots in the biplot represent the countries and their location within the plot provides a clue on the employment profile of that country in 2000-2015. For example, countries in which a high percentage of the population was employed in the agricultural sector are located towards the agricultural sector cluster. Based on this it is possible to identify countries which were the employment leaders in the agricultural sector. This is a simple application of the interpretation of the directions as discussed with the reconstitution formula. However, the clusters for industry and services lie very close to each other in the 2D projection and it is hard to discern whether there are countries specifically associated to either of the two categories.

Finding out which countries are the leaders in, say, the services sector is made possible by the respective Association Plot (Fig. 2a). In the figure one can clearly see that the service category (represented by yellow crosses) is separated from the other employment clusters (black crosses). The yellow crosses are in turn divided into two groups, which correspond to the male and female data. The countries with the highest percentage of population employed in the services are located towards the right part of the plot. Starting from right, the leaders are: Hong Kong, Luxembourg, Guam, and Kuwait followed by Macao, Saudi Arabia, Brunei Darussalam, Singapore, Netherlands and the United Kingdom. Representative barplots for some of these contries are given in Fig. 2b-e. The countries with the lowest proportion of the population in services are on the opposite side of the Association Plot (Burundi, Ruanda, Central African Republic, Niger and Rwanda). The Association Plot visualizes this information while it would be invisible in the low-dimensional projection. The Association Plot for the industry cluster (data not shown) actually indicates a lack of countries exclusively associated to industry.

**Fig. 2:**
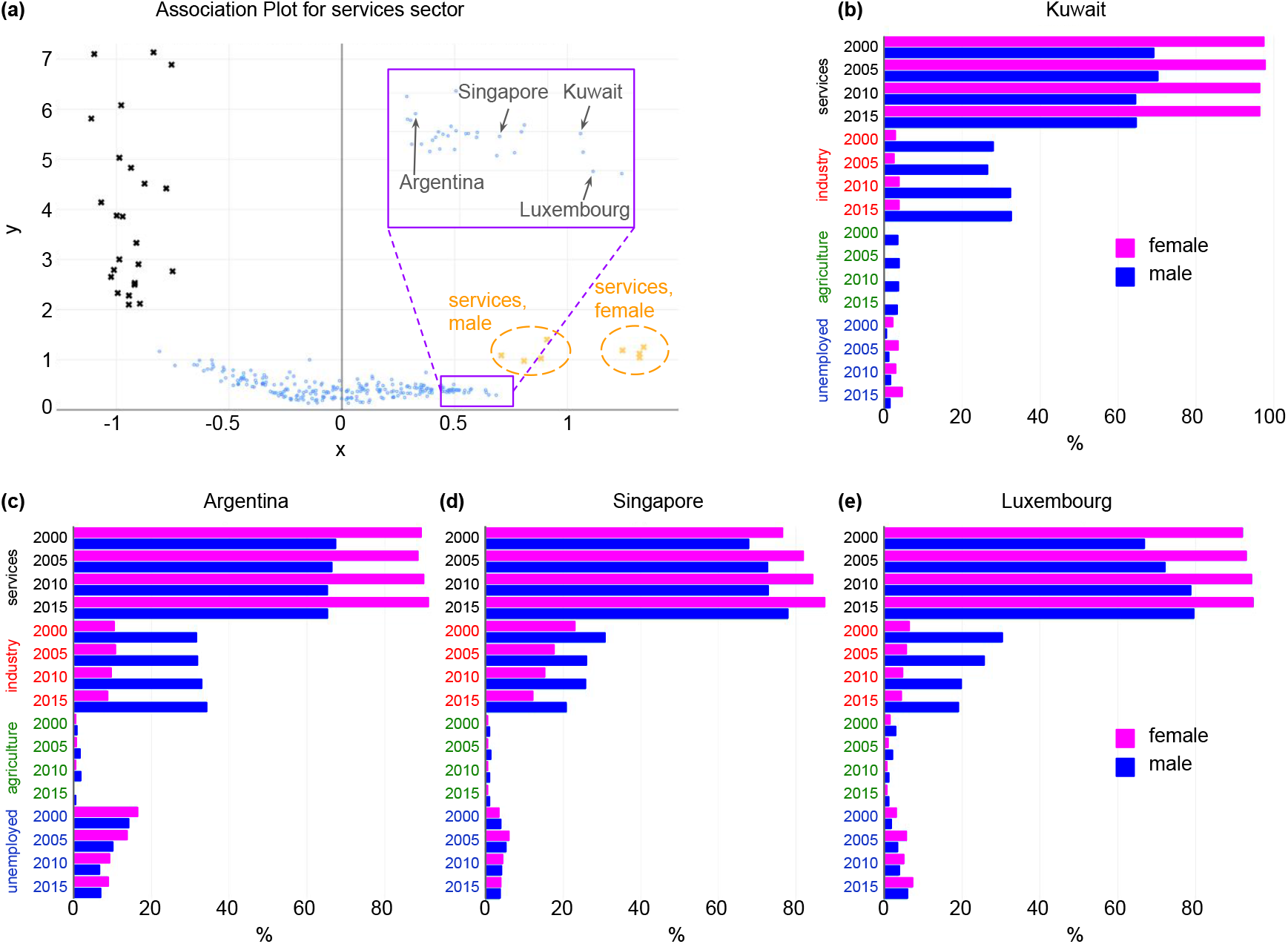
Association Plot for services sector. (**a**) The Association Plot was generated from the employment data using eight CA dimensions. The countries with the highest percentage of population employed in the services sector (yellow crosses) are located towards the right part of the plot. The names of four example leader countries are shown. The countries with the lowest proportion of population employed in services are located towards the left part of the plot, in the direction of other employment sectors (black crosses). (**b-e**) Employment profiles of the example leaders in services sector: (b) Kuwait, (c) Argentina, (d) Singapore and (e) Luxembourg. The presented barplots illustrate the percentages of female and male population in given country employed in different sectors in the years 2000-2015.

## 6. Dimensions: Noise reduction rather than projection into 2 or 3 dimensions

Traditionally, both PCA and CA are performed with the goal of depicting the information either in the plane or, possibly, in three dimensions. Where this implies a large loss of information, the SVD can still provide for noise reduction by canceling the dimensions that belong to the small singular values. This is common practice in the analysis of large data sets. Employing this mechanism implies a projection into a still high-dimensional subspace, which cannot be visualized but at least maintains the relevant information in the data.

For our purposes, we adopt this procedure for noise-reduction in the CA-space, i.e. before computing Association Plots. The estimation of the number of dimensions that should be retained, can be based on e.g. the number of clusters in the investigated data, or can be done computationally be analyzing the scree plot, i.e. the plot showing the sorted singular values from largest to smallest. We rely on the “elbow rule” which is based on a scree plot for randomized data. The dimension is read off from the point where the scree plot of the actual data enters the band of sorted randomized singular values (Ciampi *and others*, 2005).

When data happen to fall nicely into clusters, the number of dimensions to be retained should roughly reflect the number of clusters in the data. For example, the Association Plot in the employment example above was computed for eight dimensions. However, many papers have been written on the problem of selection of the right number of singular values (see the literature on spectral clustering, e.g., Zelnik-Manor and Perona, 2005; Von Luxburg, 2007). Luckily, visual inspection shows that the Association Plot is fairly robust with respect to the precise choice of dimension for the computation. An example is shown in Supplementary Fig. S1, where Association Plots were calculated for the GTEx example (see below) based on 37, 96, or 225 dimensions kept from the CA. These numbers were obtained using three different approaches for selecting optimal number of dimensions (Ciampi *and others*, 2005; Greenacre and Blasius, 1994). Inspection of the cluster-specific gene sets shows large overlaps across these choices.

## 7. Scoring of visual patterns

To obtain a better understanding of the visual patterns we observe in the Association Plot we study random data. For CA alone, randomized data would lead to a dense, roughly ellipsoidal cloud of gene-points around the origin. For a random Association Plot, however, it does not suffice to randomize the data, but one also needs to define a random cluster of conditions to orient the Association Plot in the direction of the centroid. Taken together, we first randomize a given data set by permuting the rows of the data matrix following Tusher *and others*, 2001. Next we select a number of conditions to form a random cluster. The centroid of this random cluster will determine an arbitrary direction in space. Additionally, we have observed that the cardinality of the cluster also influences the appearance of a random Association Plot. This seems to be due to the angle between the centroid and the cluster members depending on the size of a cluster.

Fig. 3a shows such an Association Plot generated from randomized data, where a “pseudo”-cluster of 600 conditions was selected for plotting the Association Plot. In fact, the randomized data in this example come from the example data set (GTEx) discussed below and the number of cluster members corresponds to the cardinality of the heart cluster. One observes a V-shaped cloud of points, which is the Association Plot’s view on the dense cloud of points around the origin in CA-space. The width of the V provides information as to the area occupied by chance and to the right of which relevant information may start. Thus, we aim to determine the angle of the ray delineating the V towards the right. The empirical distribution of the number of points falling to the right of the delineating line is shown in Fig. 3b. It allows for the choice of a threshold on the angle *α*. We have chosen the line with degree 68.8° which delineates 1% of the points to the right of the V.

**Fig. 3:**
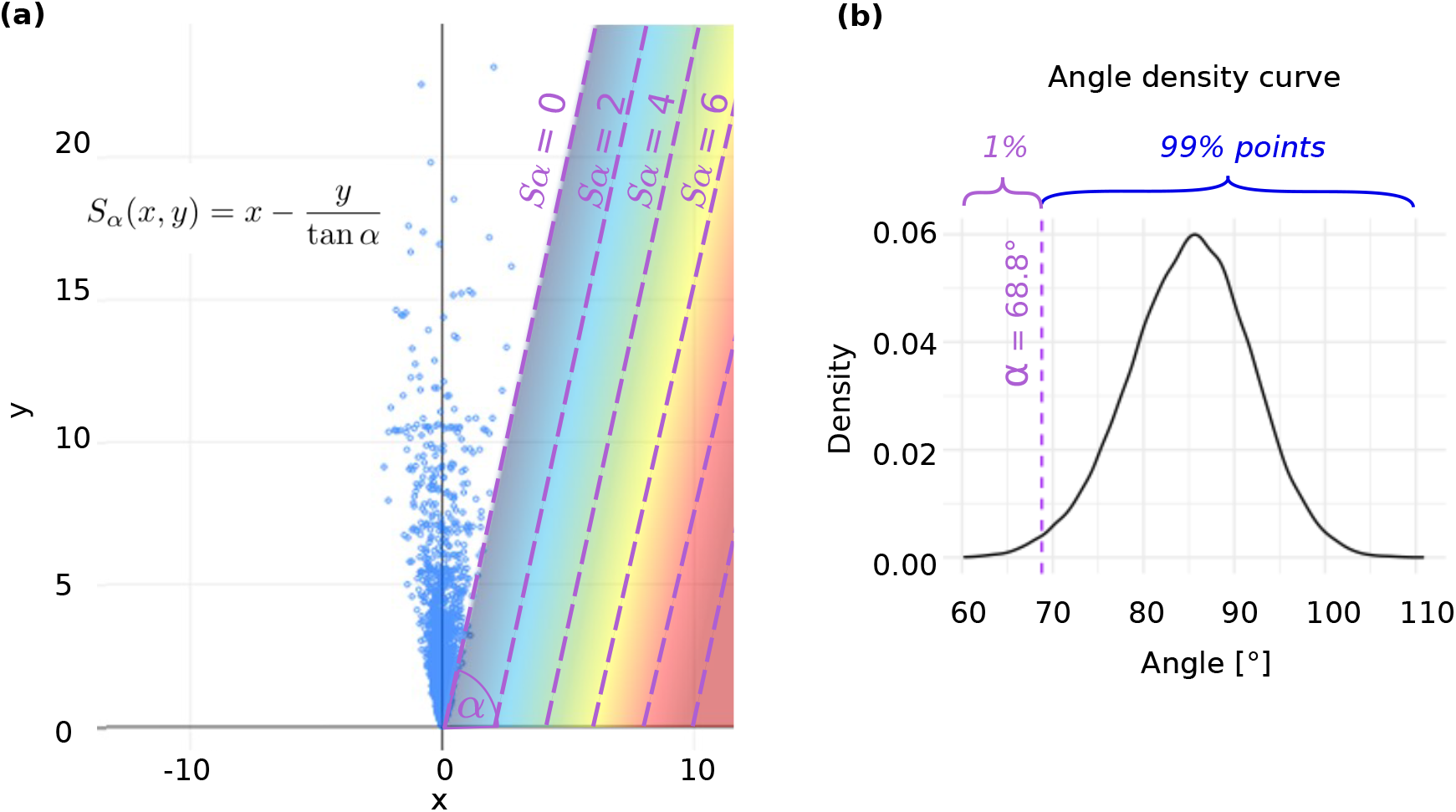
Scoring system of candidate genes. (**a**) Association Plot generated from randomized GTEx data for a “pseudo”-cluster comprising 600 conditions. For each point the angle between a given point and the x-axis was calculated. 1% of points with the lowest angle determines the *a* threshold, which will be further used for calculating the gene scores *S_α_* in the original data. (**b**) Distribution of the angles between points from (a) and the x-axis. In this example the threshold of 1% resulted in *a* = 68.8°.

Based on the angle thus determined we propose the following heuristic choice of a scoring function *S_α_(x,y)* for an individual point (*x,y*) in the Association Plot:

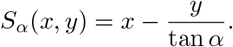

*S_α_* is 0 along the delineating line of degree *α* which the simulation yielded and which in Fig. 3a is annotated with “ *S_α_* = 0”. The scoring function *S_α_* is designed such that parallels to this line constitute level lines of increasing *S_α_* as they shift to the right (see Fig. 3a). This serves the purpose of giving higher scores to points further towards the right, while at the same time decreasing the scores as one moves upward. With this choice one adds additional power of distinguishing among genes which otherwise would have the same association value with respect to a cluster. The scoring function can, e.g., be used to rank the genes based on an Association Plot. The data example below will provide an example illustrating and supporting this choice of the scoring function *S_α_*.

## 8. A biological application: The GTEx data

To demonstrate the utility of the Association Plots in finding cluster-specific genes we present an application of our method to a large biological data set: the Genotype-Tissue Expression (“GTEx”) data (Carithers *and others*, 2015). This data set comprises 11,688 gene expression samples (matrix columns) generated by RNA-seq from 30 human tissues collected from different donors (see Methods for GTEx data processing). Each column contains one of many replicates for each tissue. The tissue information provides a natural clustering of the columns and we interpret each tissue as one cluster. The rows of the matrix correspond to the genes whose expression levels were measured, of which we use the 5,000 genes with the largest variance. Altogether the matrix has a size of 5000 x 11,688, and the question we address is “Which genes are associated to a certain tissue?”. In biology, this corresponds to the search for so-called marker genes for a tissue.

For demonstration purposes we first conducted correspondence analysis and projected the data into a three-dimensional subspace. The three dimensions of this projection together explain only ca. 24% of the inertia in this data set. The plot is clearly organized around the three directions for the tissues: pancreas, blood, and pituitary gland (Fig. 4a). Thus, the Association Plot for pancreas (Fig. 4b) contains no surprise: Many genes point to the right, in the direction of the pancreas centroid. We provide a zoom into the right tail of the plot. This clearly shows a set of pancreas-specific genes, of which we colored in red the known pancreas marker genes as determined by the Human Protein Atlas (Uhlén *and others*, 2015).

**Fig. 4:**
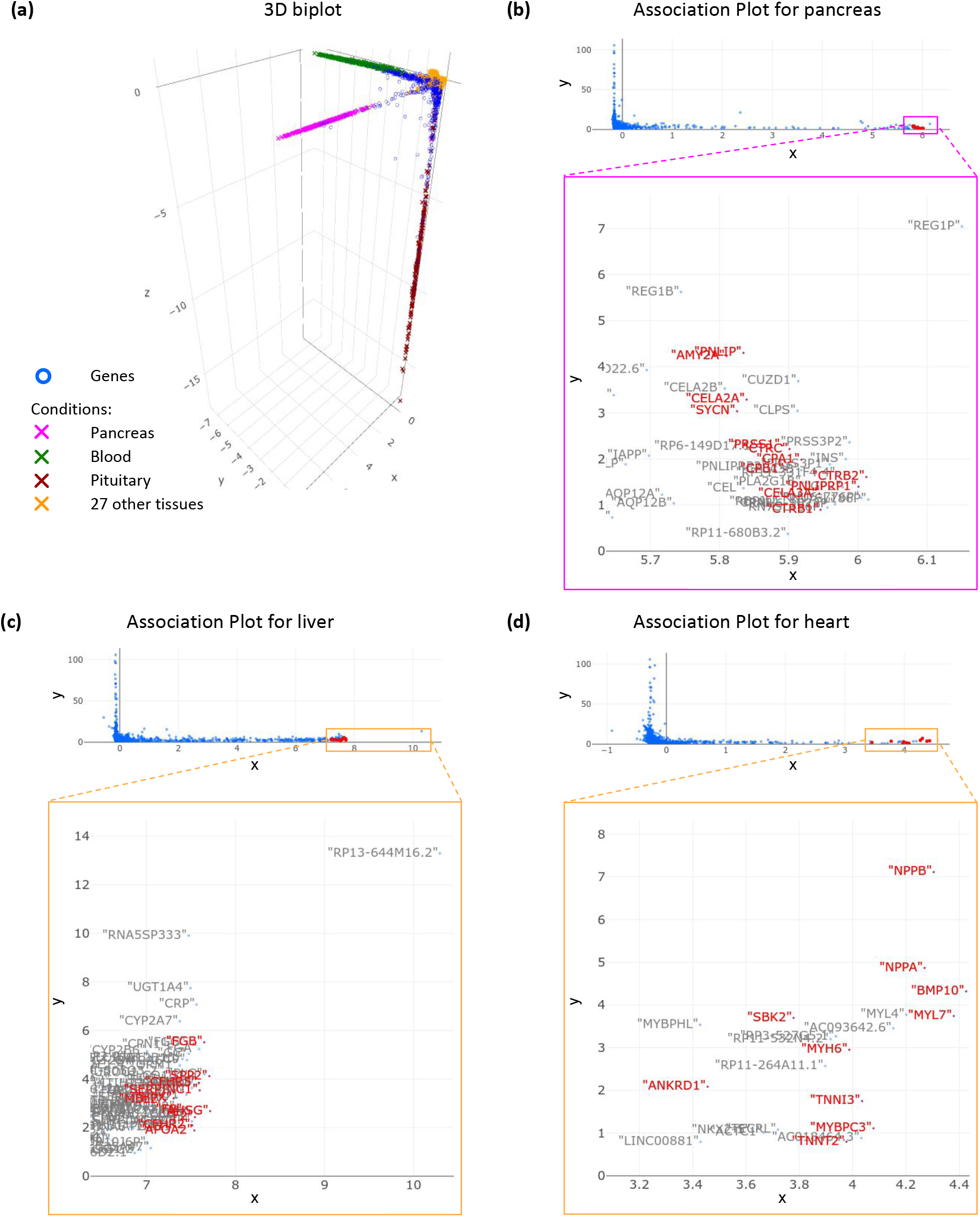
Applying CA to GTEx data. (**a**), Classical CA projection into 3D subspace, which allows discerning three different tissues (pancreas, blood, pituitary). Other tissues remain lumped in the centre of the observed structure. Association Plots enable obtaining further tissue-specific information. **b-d,** The Association Plots were generated based on the first 96 CA dimensions and can be used for delineating pancreas- (**b**), liver- (**c**) and heart-specific (**d**) genes. Such genes are located in the right bottom part of the plot. Red color indicates marker genes from Human Protein Atlas for a given tissue. The presented Association Plots serve as examples. Each tissue can be inspected separately and respective marker genes can be visualized.

The real challenge, however, lies in the invisibility of the remaining 27 tissues, which cannot be distinguished from each other since they form a dense cloud around the origin of the coordinate system. Consequently, it is also impossible in the 3D-projection to identify marker genes for these tissues. This is where the Association Plots come in and enable extracting further tissue-specific information. As an example, Fig. 4c depicts the Association Plot generated for the liver samples. Again, a zoom into the right tail of the plot shows numerous genes, a subset of which are known as liver-specific marker genes as given by the Human Protein Atlas (Uhlén *and others*, 2015). One can in principle go through the tissues in this manner and as another representative example heart-specific genes are shown in the Association Plot in Fig. 4d.

The GTEx data example shall also serve to exemplify the score *S_α_*. Fig. 5a shows the heart-specific genes colored by the *S_α_* value. The angle *α* is taken from the simulated data, which are exactly the ones shown above in Fig. 3. The ranking of genes provided by the color code is a rather intuitive ranking. Some examples of the actual distribution of gene expression values over the tissues are shown in Fig. 5b and provide an intuition of the associations depicted in these ranked Association Plots.

**Fig. 5:**
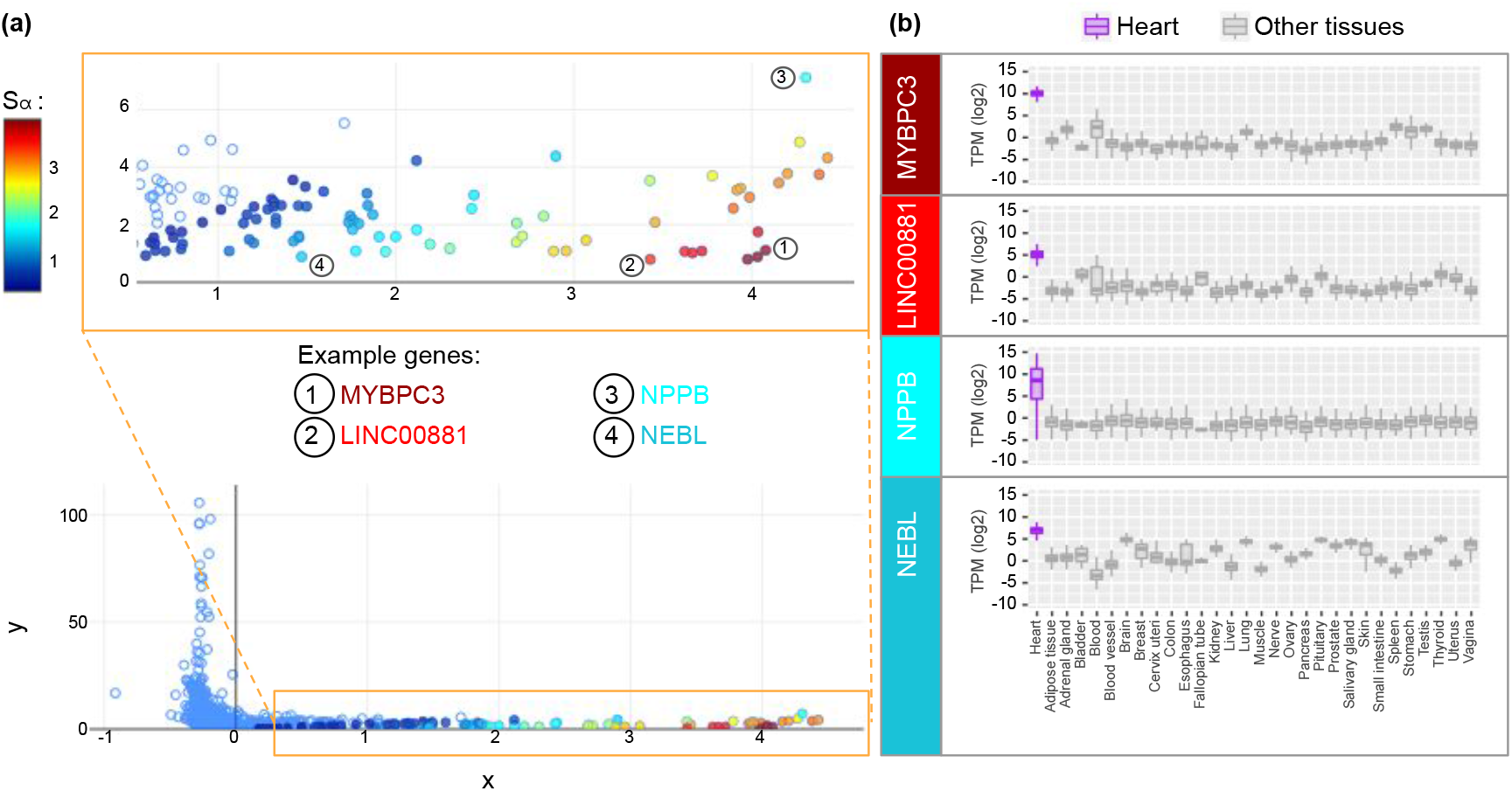
Heart-specific genes colored by their *S_α_* values. (**a**) Association Plot was generated for heart samples from GTEx data using 96 CA dimensions. The color of a gene (circle) refers to its *S_α_* value. The higher the *S_α_* value, the higher the specificity of the given gene for heart samples. (**b**) Boxplots illustrating the expression of four example genes (MYBPC3, LINC00881, NPPB, and NEBL) across 30 tissues. The boxplots generated for genes with high *S_α_* values clearly show their upregulation in the heart tissue in comparison to other tissues. When comparing the expression profiles of the four example genes with higher *S_α_* values, MYBPC3 shows the highest upregulation in the heart tissue, whereas NEBL the lowest.

The Association Plots for the GTEx data were computed based on the first 96 dimensions of the CA-space, as determined by the elbow rule. Supplementary Fig. S1 shows the very similar images for 37, 96, and 225 dimensions.

## 9. Discussion

Motivated mostly by biological questions, we developed and presented here a novel visualization method, Association Plots, to study cluster-specific genes in a contingency table. As a generic data visualization method, Association Plots do not make particular distributional assumptions as would be commonly made in the field of gene expression. For example, the widely used DE-Seq2 (Love *and others*, 2014) program for determining differentially expressed genes assumes that data follow a negative binomial distribution. In this context we see Association Plots strictly as a visualization tool and do not claim the same statistical rigor as the specialized methods.

Generally, data sets organized in form of a matrix or contingency table are ubiquitous, and today their size and complexity can be intimidating. Cluster information is also frequently available, be it a natural clustering stemming from the nature of the data, or a computed one. With large size of a data set, a traditional exploratory visualization method like, e.g., PCA, may provide little help, whereas Association Plots offer the capability to visualize and interact with the sets of rows (or genes) that are associated to a cluster of columns (or conditions). Association Plots can provide intuitive visualization in a manner only dependent on the clusters to be studied but independent of the size of the data set. Dimension reduction is applied merely for the purpose of noise reduction and not to a degree where the visualization would cancel significant amounts of information from the data.

At the heart of our method lies the geometry of the correspondence analysis biplot, where the direction from the origin towards a column/condition allows for an interpretation in terms of row-column association. This gives rise to the representation of rows by orthogonal distance to this direction vector. What is visualized is essentially a cone from the origin centered on a column or a cluster centroid. This makes the geometry of correspondence analysis a necessary prerequisite for Association Plots.

An aspect often overlooked in data analysis methods is whether a method will indicate that its assumptions are not met. Association Plots can also reveal that a given cluster of conditions is in fact not a coherent cluster. In this case the typical structure of genes pointing in the direction of the cluster centroid in CA-space will be dispersed, and, as a consequence, the right tail in the Association Plot generated for this cluster will be short and not clearly visible anymore. Flagging the violation of a clustering assumptions is a desirable feature of our method.

We believe that the viewpoint on data analysis described opens up many further interesting questions. Most prominently, we plan to study the connection between Association Plots and biclustering (Tanay *and others*, 2002; Pontes *and others*, 2015). Also the connection to spectral clustering (Zelnik-Manor and Perona, 2005; Von Luxburg, 2007) mentioned above seems worth pursuing.

## 10. Data and Methods

### 10.1 GTEx data

RNA-seq data of postmortem non-disease human tissues was retrieved from GTEx Portal (Carithers *and others*, 2015). The files “GTEx\_Analysis\_2016-01-15\_v7\_RNASeQCv1.1.8\ _gene\_tpm.gct.gz” containing gene TPM values and “ GTEx\_v7\_Annotations\_SampleAttributesDS. txt” containing sample annotations (both files available from https://gtexportal.org/home/ datasets) were downloaded on 03.07.2019. A detailed description of data processing procedures is available from https://gtexportal.org/home/documentationPage.

Correspondence analysis was computed using 5,000 genes with the highest expression variance across all 11,688 samples. For generating the Association Plots in Fig. 4 the first 96 CA dimensions (number obtained using the elbow rule) were considered.

### 10.2 Human tissue marker genes

For validation of the Association Plots generated for GTEx data, the lists of genes with the highest levels of enriched expression in liver, heart, and pancreas were obtained on 03-18-2019 from the Human Protein Atlas (Uhlén *and others*, 2015), available from http://v18.proteinatlas.org.

### 10.3. Code availability

The code to produce Association Plots is written (mostly) in *R* using shiny library (Chang *and others*, 2019) and is available from the GitHub repository https://github.com/elagralinska/ APL. Only the SVD routine (torch.svd) is taken from Python3 torch package and gets called from the *R* code.

## Acknowledgments

We thank our colleagues Robert Schöpflin, Peter Holderrieth and Verena Heinrich for providing feedback and criticism on the developing manuscript.

---

**Fig. S1:**
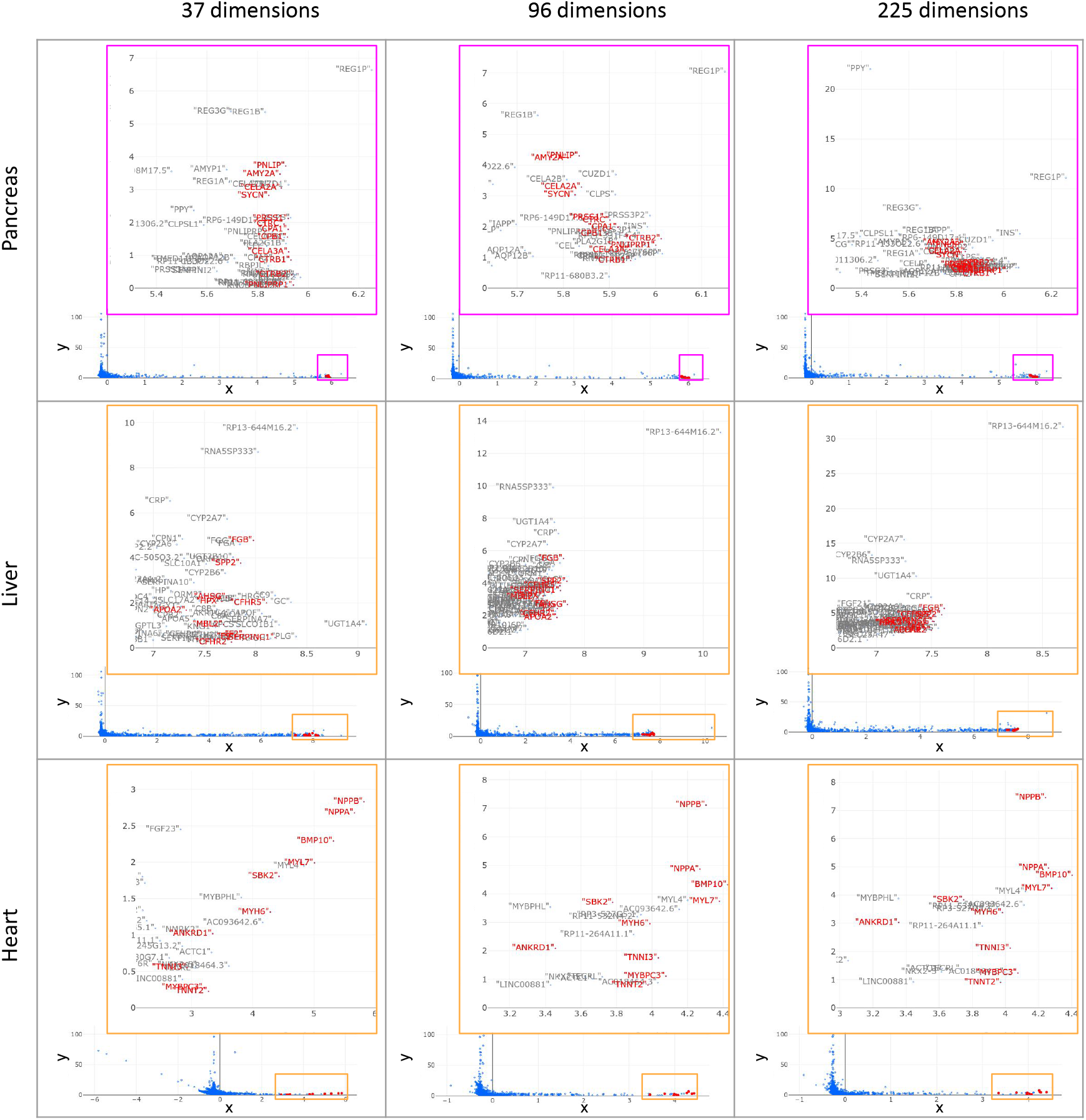
Comparison of Association Plots generated for three tissues (pancreas, liver, heart) using three different numbers of dimensions. The numbers of dimensions (37, 96 and 225) were calculated using three different approaches (keeping only those dimensions that explain more inertia than one dimension on average; keeping the minimum number of dimensions that account for more than 80% of inertia in total; elbow rule (Ciampi *and others*, 2005)). For each tissue the cluster-specific gene sets obtained using the three different approaches show large overlaps.

## References

Carithers, Latarsha J, Ardlie, Kristin, Barcus, Mary, Branton, Philip A, Britton, Angela, Buia, Stephen A, Compton, Carolyn C, DeLuca, David S, Peter-Demchok, Joanne, Gelfand, Ellen T and others. (2015). A novel approach to high-quality postmortem tissue procurement: the gtex project. Biopreservation and biobanking 13(5), 311–319.

Chang, Winston, Cheng, Joe, Allaire, JJ, Xie, Yihui and McPherson, Jonathan. (2019). shiny: Web Application Framework for R. R package version 1.3.2.

Ciampi, Antonio, González Marcos, Ana and Castejón Limas, Manuel. (2005). Correspondence analysis and 2-way clustering. SORT 29(1).

Greenacre, Michael. (2017). Correspondence analysis in practice. Chapman and Hall/CRC, Boca Raton (FL).

Greenacre, Michael J. (1984). Theory and applications of correspondence analysis. Academic Press, London.

Greenacre, Michael J and Blasius, Jörg. (1994). Correspondence analysis in the social sciences: Recent developments and applications. Academic Press, London.

International Labour Organization. (2020). Ilostat database [database]. Available from World Development Indicators: http://data.worldbank.org/indicator/SL.SRV.EMPL.MA.ZS, http://data.worldbank.org/indicator/SL.SRV.EMPL.FE.ZS, http://data.worldbank.org/indicator/SL.IND.EMPL.MA.ZS, http://data.worldbank.org/indicator/SL.IND.EMPL.FE.ZS, http://data.worldbank.org/indicator/SL.AGR.EMPL.MA.ZS, http://data.worldbank.org/indicator/SL.AGR.EMPL.FE.ZS, http://data.worldbank.org/indicator/SL.UEM.TOTL.MA.ZS, http://data.worldbank.org/indicator/SL.UEM.TOTL.FE.ZS.

Love, Michael I, Huber, Wolfgang and Anders, Simon. (2014). Moderated estimation of fold change and dispersion for rna-seq data with deseq2. Genome biology 15(12), 550.

Pontes, Beatriz, Giráldez, Raúl and Aguilar-Ruiz, Jesus S. (2015). Biclustering on expression data: A review. Journal of biomedical informatics 57, 163–180.

Tanay, Amos, Sharan, Roded and Shamir, Ron. (2002). Discovering statistically significant biclusters in gene expression data. Bioinformatics 18(suppl_1), S136–S144.

Tusher, Virginia Goss, Tibshirani, Robert and Chu, Gilbert. (2001). Significance analysis of microarrays applied to the ionizing radiation response. Proceedings of the National Academy of Sciences 98(9), 5116–5121.

Uhlén, Mathias, Fagerberg, Linn, Hallström, Bjorn M, Lindskog, Cecilia, Oksvold, Per, Mardinoglu, Adil, Sivertsson, Asa, Kampf, Caroline, Sjöstedt, Evelina, Asplund, Anna and others. (2015). Tissue-based map of the human proteome. Science 347(6220), 1260419.

Von Luxburg, Ulrike. (2007). A tutorial on spectral clustering. Statistics and computing 17(4), 395–416.

Wang, Zhong, Gerstein, Mark and Snyder, Michael. (2009). Rna-seq: a revolutionary tool for transcriptomics. Nature reviews genetics 10(1), 57–63.

Zelnik-Manor, Lihi and Perona, Pietro. (2005). Self-tuning spectral clustering. In: Advances in neural information processing systems. pp. 1601–1608.

